# Plasticity in medaka gonadotropes via cell proliferation and phenotypic conversion

**DOI:** 10.1101/757682

**Authors:** Romain Fontaine, Eirill Ager-Wick, Kjetil Hodne, Finn-Arne Weltzien

## Abstract

Follicle stimulating hormone (Fsh) and luteinizing hormone (Lh) produced by the gonadotropes, play a major role in control of reproduction. Contrary to mammals and birds, Lh and Fsh are mostly produced by two separate cell types in teleost. Here, we investigated gonadotrope plasticity, using transgenic lines of medaka (*Oryzias latipes*) where DsRed2 and hrGfpII are under the control of fshb and lhb promotors respectively. We found that Fsh cells first appear in the pituitary at 8 dpf. Similar to in Lh cells, Fsh cells show hyperplasia from juvenile to adult stages. Hyperplasia is stimulated by estradiol exposure. Both Fsh and Lh cells show hypertrophy during puberty with similar morphology. They also share similar behavior, using their cellular extensions to make networks. We observed bi-hormonal gonadotropes in juvenile and adult fish but not during larval stage where only mono-hormonal cells are observed, suggesting the existence of phenotypic conversion between Fsh and Lh in later stages. This is demonstrated in cell culture, where some Fsh start to produce *lhb*, a phenomenon enhanced by gonadotropin releasing hormone (Gnrh) stimulation. We have previously shown that medaka Fsh cells lack Gnrh receptors, but here we show that with time in culture, some Fsh cells start responding to Gnrh, while *fshb* mRNA levels are significantly reduced, both suggestive of phenotypic change. All together, these results reveal high plasticity of gonadotropes due to both estradiol sensitive proliferation and Gnrh promoted phenotypic conversion, and also shows that gonadotropes lose part of their identity when kept in cell culture.

## INTRODUCTION

Gonadotropes are key players in the control of the reproductive function as part of the Brain-Pituitary-Gonad axis (Harris 1951; Weltzien, et al. 2004). Located in the anterior part of the pituitary, they produce the two gonadotropins: follicle-stimulating hormone (Fsh) and luteinizing hormone (Lh) (Weltzien, et al. 2014). Fsh and Lh are mostly produced by the same cell in mammals (Nakane 1970), while the opposite occurs in teleost fish, where Fsh and Lh are produced by two different cell types (Kanda, et al. 2011; Nozaki, et al. 1990; Schmitz, et al. 2005; Weltzien, et al. 2014). Therefore, teleosts seem ideal models to study the development and the different properties of Fsh and Lh cells, as well as the differential regulation of Fsh and Lh synthesis and release (Weltzien, et al. 2014; Yaron, et al. 2003).

However, despite the general understanding of one hormone one cell type in teleosts, several observations have challenged this hypothesis. Indeed, gonadotropes producing both gonadotropins were found in several teleost species (e.g. Mediterranean yellowtail (Hernandez, et al. 2002), zebrafish, tilapia (Golan, et al. 2014) and European hake (Candelma, et al. 2017)). On the other hand, gonadotropes expressing only one hormone were described in mammals (Childs 1983; Childs, et al. 1982). Previous publications have pointed out the fact that Fsh and Lh share the same developmental basis in fish, similar to what is found in mammals (Weltzien, et al. 2014) suggesting that Fsh and Lh cells may not be so different from each other in fish, and more similar to the mammalian gonadotropes than we perhaps have anticipated.

Medaka is a powerful teleost model for which several tools have been developed to study its genetics and development (Shima and Mitani 2004; Wittbrodt, et al. 2002). The recent development by our team of two transgenic lines, where DsRed2 and hrGfpII reporter proteins synthesis are controlled by the endogenous medaka *fshb* and *lhb* promotors respectively, enables the study of the gonadotrope cells in more detail (Hildahl, et al. 2012; Hodne, et al. 2019).

Previously several studies conducted on Lh cells in medaka have explored and investigated basic parameters including morphology, ontogeny and regulation of Lh cells. In medaka, Lh cells have been found to participate in the plasticity of the pituitary during puberty through hypertrophy and estrogen-sensitive hyperplasia during puberty (Fontaine, et al. 2019). Lh cells have also been shown to make neuron-like projections allowing homotypic networks (Grønlien, et al. 2019), to express gnrh receptors (gnrhr), and to respond to gnrh stimuli by increasing their action potential frequency and intracellular calcium concentration (Hodne, et al. 2019; Strandabo, et al. 2013). However, very little is known about Fsh cells and if and how they contribute to pituitary plasticity during puberty. Therefore, using the recently developed transgenic lines where Fsh and Lh cells can be identified, we investigated gonadotrope plasticity in the medaka pituitary, examining both proliferation and phenotypic plasticity. In addition, we investigated the presence and the origin of bi-homonal (Fsh and Lh) cells in medaka.

## MATERIALS AND METHODS

### Animal maintenance

Wild-type (WT, d-rR strain), transgenic tg(*lhb*-hrGfpII) (Hildahl, et al. 2012), tg(*fshb*-DsRed2) and double transgenic tg(*lhb*-hrGfpII/*fshb*-DsRed2) (Hodne, et al. 2019) medaka (*Oryzias latipes*) were maintained at 28°C on a 14/10 hr light/dark cycle in a re-circulating system with reverse osmosis dosed-salt water (pH 7.6 and conductivity of 800μs). Fish were fed three times a day, once with live brine shrimp and twice with dry feed (Gemma, Skretting, UK). Experiments were performed according to the recommendations of the care and welfare of research animals at the Norwegian University of Life Sciences, and under the supervision of authorized investigators. Specifically, the Bromodeoxyuridine (BrdU) experiments were approved by the Norwegian Food Safety Authorities (FOTS ID 8596).

### Primary pituitary dispersed cell cultures

Cell cultures were prepared as described in detail in (Ager-Wick, et al. 2018). For measuring the volume of DsRed2 and hrGfp-II expressing cells, 4 cell cultures were prepared either from 15 adult or due to their smaller size, 25 juvenile pituitaries of tg(*lhb*-hrGfpII/*fshb*-DsRed2) animals from each sex. For quantification of mRNA levels at different time points, cell cultures were prepared by dissociating cells from 25 adult tg(*lhb*-hrGfpII) females. Cells were then plated in 3 different wells (each corresponding to a different sampling time point) in a 48 wells plastic plate (Sarstedt, Germany) coated with poly-L-lysine (Sigma, Norway), prepared in a laminar flow hood by adding 50 μl poly-D-lysine, leaving for 1 min before removing the liquid, washing in 500 μl MQ water and leaving the coated wells to dry in UV-light for approximately 30 minutes.

For investigation of phenotypic conversion, 6 cell cultures from males and 4 from females tg(*lhb*-hrGfpII/*fshb*-DsRed2) were prepared. 2 cell cultures from each sex were treated 4 hours after being plated by adding Gnrh1 (concentration 10^−6^ M) into the medium. Time lapse was recorded as described below for 3 days.

### qPCR

i) fshb mRNA was quantified during development using WT medaka as described in (Hildahl, et al. 2012). Briefly, a LightCycler 480 Real-Time PCR system (Roche, Mannheim, Germany), with SYBR Green (Roche) was used. Pooles of synchronized embryos (see table 2 in (Hildahl, et al. 2012)) were collected in RNAlater for RNA isolation and cDNA synthesis. ii) *gnrhr1b*, *gnrhr2a*, *gnrhr2b*, *lhb* and *fshb* mRNA were quantified from cell cultures at 3 different time points: 1 hour, 24 hours and 72 hours after plating the dissociated cells. Cells where mechanically detached from the plate by scraping the cells using the pipette in 300 ul of Trizol and further submitted to phenol-chloroform RNA extraction using GlycoBlue (Invitrogen, California, USA) as carrier. Experiments were performed in quadruplicate and triplicate respectively, for proper statistical analysis. Primers for *gnrhr1b*, *gnrhr2a*, *gnrhr2b*, *lhb, fshb* and 4 reference genes (β*-actin*, *glyceraldehyde 3-phosphate dehydrogenase* (*gapdh) ribosomal protein L7* (*rpl7*), *16s* and *18s ribosomal RNA* (*16s* and *18s*)) were designed to span exon–exon boundaries based on in silico analysis of the medaka genome to avoid detection of genomic DNA (gDNA) (Table 1). *16s* expression was found to be the most stable across larval development and the combination of *gapdh*, *rpl7* and *18s* was found to be the most stable across time in cell culture according to BestKeeper software (Pfaffl, et al. 2004), and thus used to normalize the expression analysis, using an efficiency-corrected relative quantification method (Weltzien, et al. 2005).

**Table 1:**
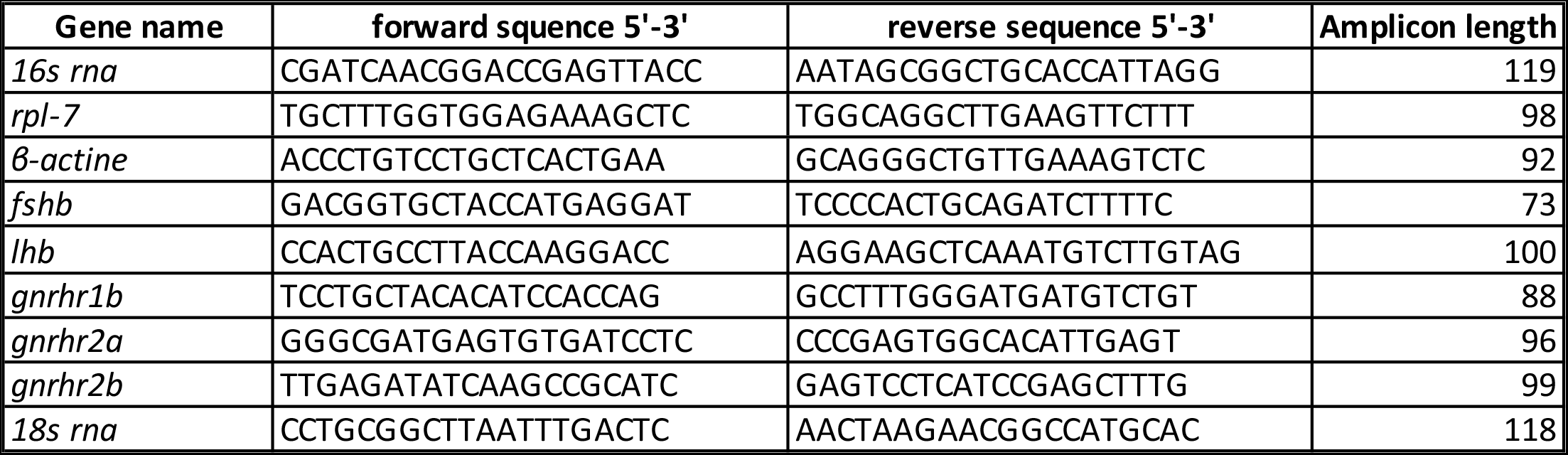
List of primers used for qPCR.

### Steroid treatments and BrdU Incubation

To study effects of sexual steroids on Lh and Fsh cell proliferation, 3 groups of transgenic tg(*lhb*-hrGfpII) adult fish (6 females and 6 males) were incubated for 6 days in system water containing 100 μg/L of either 17β-estradiol, testosterone or 11-ketotestosterone (Sigma; diluted 1:10^5^ in 96 % ethanol). Control fish (6 of each sex) were incubated for 6 days with diluent only. The experiment was repeated once. Immediately after steroid treatment, the fish were treated with 1 mM BrdU (Sigma) diluted in water with 15 % DMSO for 8 h. Fish were then sacrificed, brain and pituitary were collected and fixed in 4 % paraformaldehyde overnight, and gradually dehydrated and stored in 100 % MetOH until use.

### Immunofluorescence

Tissues were labeled for BrdU, PCNA as well as for Fshβ with immunofluorescence (IF), as previously described (Burow, et al. 2019; Fontaine, et al. 2013; Fontaine, et al. 2019). Briefly, IF was performed on free-floating sections obtained after the tissues were included in 3% agarose and parasagittaly sectioned (60 μm) with a vibratom (Leica). Because the fluorescence of the endogenous DsRed2 is quenched with the epitope retrieval treatments required for BrdU and PCNA staining, tg(*lhb*-hrGfpII) animals were used and IF for Fshβ, with a custom-made polyclonal rabbit anti-medakaFshβ (1:500 (Burow, et al. 2019)) was performed. Nuclei were stained with DAPI (1:1000; 4’,6-diamidino-2-phenylindole dihydrochloride; Sigma).

### Imaging

For imaging of the tg(*lhb*-hrGfpII/*fshb*-DsRed2) line during ontogeny (8-10 unsexed fish per stage) or for investigating the presence of bi-hormonal cells (12 unsexed fish per stage), no treatments where needed and endogenous hrGfpII together with DsRed2 were directly visualized. For all, vibratome slices were mounted between slide and coverslip with antifade mounting medium Vectashield (Vector, UK), and spacers were added between the slice and the coverslip when mounting whole pituitaries. Time-lapse recordings of dissociated pituitary cells were performed in a humid chamber at 26 °C with 1 % CO_2_ (Ager-Wick, et al. 2018). All confocal images were acquired using a LSM710 microscope (Zeiss, Germany) with 10X, 25X, 40X or 63X (respectively N.A. 0.3, 0.8, 1.2 or 1.4) objectives. Channels were acquired sequentially to avoid signal crossover between filters. Z-projections from confocal image stacks were obtained using Fiji software (v2.0.0 (Schindelin, et al. 2012)). 3D reconstruction was built using 3D-viewer plugin (Schmid, et al. 2010).

### Calcium imaging and Gnrh1 stimulation

Calcium imaging and Gnrh1 stimulation were performed as described in (Hodne, et al. 2019). Briefly, A total of 3 dishes of dissociated adult female tg(*fshb*-DsRed2) pituitary cells were used. Following 3 days in culture, the cells were gently washed in artificial BSA-free extracellular solution (ECS: in mM: NaCl 134, KCl 2.9, MgCl_2_ 1.2, HEPES 10, and glucose 4.5, pH 7.75 and 290 mOsm), then incubated in 5 μM Fluo4-AM dye (ThermoFisher Scientific, Massachusetts, USA) for 30 min before incubation in ECS added 0.1 % BSA for 20 min. In total, 29 cells were stimulated with Gnrh1 (10 μM dissolved in ECS with 0.1% BSA; Bachem) using puff ejection (20 kPa through a 2 MΩ glass pipette, 30-40 μm from the target cell). Cells were imaged using a sCMOS camera (optiMOS, QImaging, British Columbia, Canada) with exposure time 50 to 80 ms and sampling frequency 0.5 Hz using μManager software, v1.4 (Edelstein, et al. 2014). Relative fluorescence intensity was calculated after background subtraction as changes in fluorescence (F) divided by the average intensity of the first 15 frames (F0). Data analysis was performed using Fiji software.

### Countings and measurments

Counting of Fsh cells was performed blindly using Cell Profiler software (v2.1.0 (Carpenter, et al. 2006)) as described in (Fontaine, et al. 2019), from 8 to 9 animals from each sex and stage. Double-labeled cells (BrdU/hrGfpII or BrdU/DsRed2) after steroid and BrdU treatments were manually counted using Fiji software and cell-counter plugin. Cell volume was measured by recording Z-stacks of dissociated cells a few minutes after being plated and using Fiji software and the voxel counter plugin for 9 to 36 cells per group. The fluorescence intensity in the mean region of interest (ROI) was measured with Fiji on 5 different cells from 2 different cell cultures using 10× objective. For good clarity of the figure only 3 cells were kept.

### Statistics

Data were analyzed using GraphPad Prism (v8.0, USA) with significance set at P<0.05.

Potential differences in *fshb* mRNA levels during development, pituitary cell number or proportion and effects of sex steroids on cell proliferation were tested by one-way ANOVA followed by Tukey’s multiple comparison test. Two-way ANOVA with Tukey’s multiple comparison test was used to test for differences between mRNA levels in cell cultures sampled at different time points and between the volume of the cells in cell culture. Finally, non-parametric Mann Whitney test was used to investigate significant difference in the proportion of DsRed2 cells changing phenotype in cell culture with or without Gnrh1 stimulation.

## RESULTS

### Ontogeny of Fsh cells in the pituitary

qPCR (Figure 1A) shows that relative expression of *fshb* mRNA in the embryo starts to increase after 72 hours post fertilization (hpf; 3 days). It becomes significantly different from the early time points after 336 hpf (14 days). To investigate at which time the first Fsh cells appear, we looked at the endogenous DsRed2 (Figure 1B) fluorescence starting with adult fish, back to younger stages in the tg(*fshb*-DsRed2) line. First, we did not observe any DsRed2 cells outside of the pituitary at all studied stages. Second, we found the first cells to arise around 8 days post fertilization (dpf), with a single DsRed2 cell observed in 2 of 8 studied embryos at this stage. Third, we observed an increasing number of DsRed2 between each studied stage along development.

**Figure 1:**
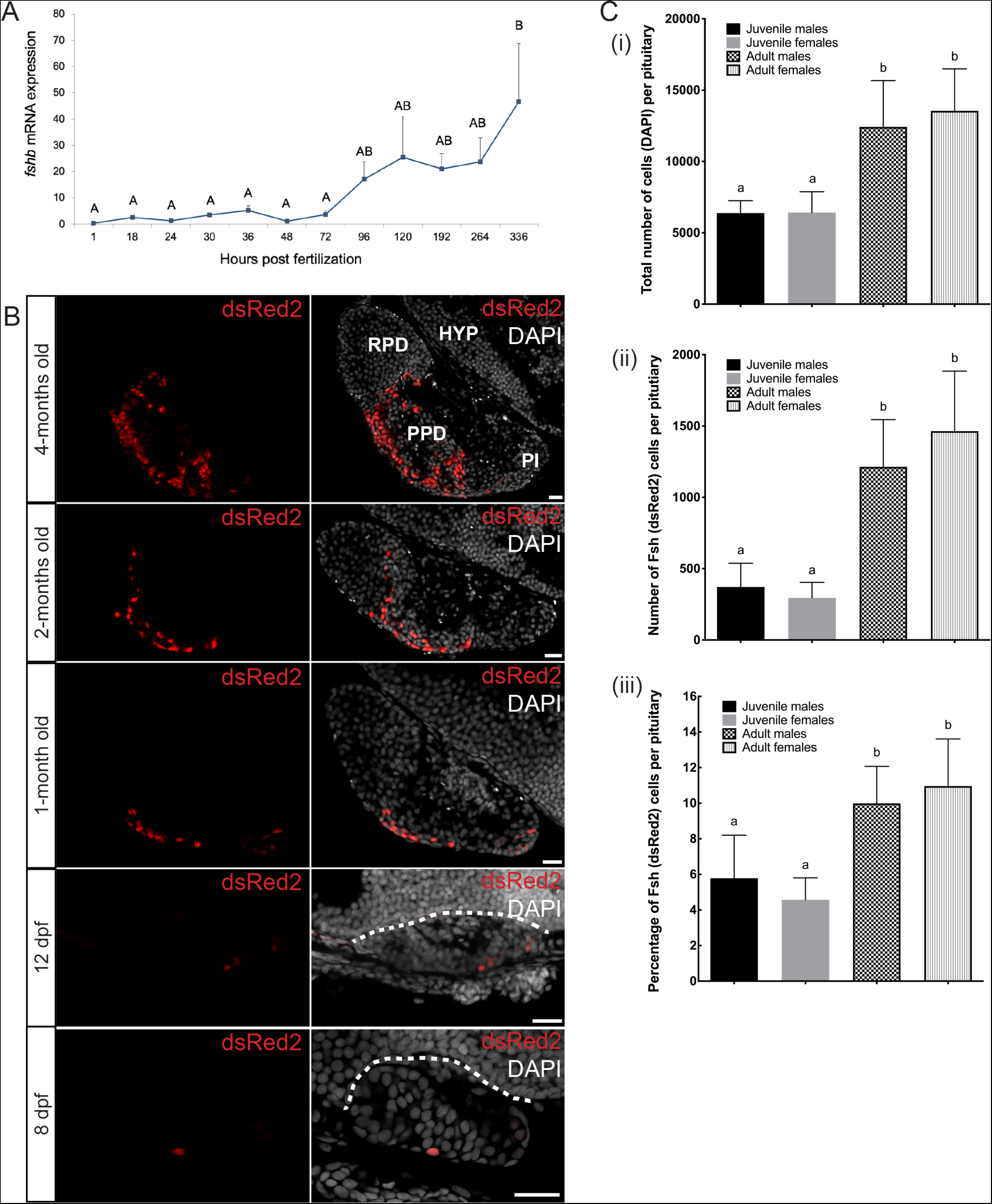
(A) Relative *fshb* mRNA expression during early development in pooled medaka larvae by quantitative polymerase chain reaction (qPCR) analysis. *fshb* gene expression was normalized to *16s* gene expression using an efficiency adjusted relative quantification method. Data are presented as mean relative expression + SEM, n=4. Relative mRNA levels were significantly different (P < 0.05) using one-way ANOVA followed by a Tukey-Kramer HSD post-hoc analysis when letters are different (A and B). (B) Ontogeny of DsRed2 producing cells in the tg(*fshb*-DsRed2) line. Parasagittal sections of the brain and the pituitary for fish from 1-month old up to 4-months old, and of the whole embryo for younger stages, without (left panels) or with nuclear (DAPI) staining (right panels). Dotted lines delimit the dorsal part of the pituitary. Scale bars: 20 μm. (C) Cell counting for the four different groups of fish: juvenile males (n = 9) and females (n = 9), and adult males (n = 8) and females (n = 9). (i) Mean (+s.d.) of the total number of cells in the pituitary. (ii) Mean (+s.d.) of the number of DsRed2 cells in the pituitary. (iii) Mean (+s.d.) of the percentage of DsRed2 cells related to the total number of cells in the pituitary. For each graph, one-way ANOVA with Tukey’s multiple comparison test revealed significant differences (P < 0.05) when letters are different (a and b).

Therefore, we counted the number of DsRed2 cells in the pituitary as well as the total number of cells using the nuclear DAPI staining, and calculated the percentage of DsRed2 cells in the pituitary (Figure 1C), in juveniles (2-month old) and adults (6-month old) in both sex. While the number of cells in the pituitary increased significantly between juvenile and adult stages, there was no significant differences between sex at any stage. The same observation was made for the number of DsRed2 cells and the percentage of DsRed2 cells in the pituitary. In adults however, there was a noticeable tendency for higher numbers of cells and DsRed2 positive cells in females as compared to in males.

### Proliferation of Fsh cells

We then looked for the origin of the new DsRed2 positive cells. IF for proliferating cell nuclear antigen (PCNA) together with Fshβ showed some cells expressing both proteins (Figure 2A-C). In addition, IF for BrdU together with Fshβ on fish incubated for 8 hours in BrdU solution revealed that some Fshβ producing cells had integrated BrdU, therefore confirming active cell division (Figure 2D-F).

**Figure 2:**
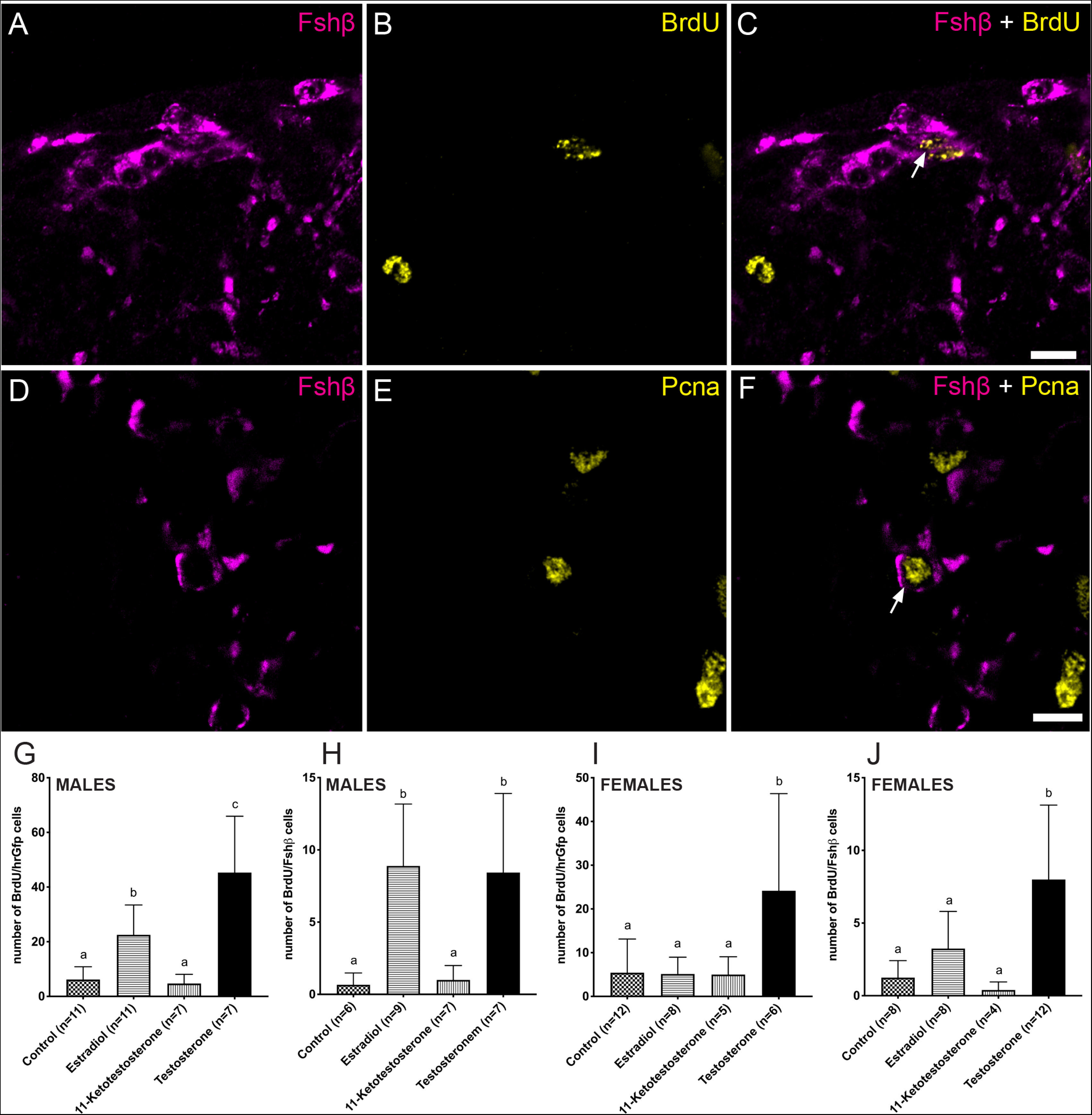
(A-C) Confocal plan images of a parasagittal section from an adult WT female medaka pituitary labeled by immunofluorescence for Fshβ (magenta) and proliferating cell nuclear antigen (PCNA; yellow). (D-F) Confocal plan images of a parasagittal section from a tg(*lhb*-hrGfpII) adult female medaka pituitary incubated in BrdU for 8 hours and labeled by immunofluorescence for Fshβ (magenta) and BrdU (yellow). Scale bars: 10 μm. (G-J) Graphics presenting the mean (+s.d.) number of double labelled cells, BrdU/hrGfpII (G,I) or BrdU/Fshb (H,J) in the pituitary from adult medaka males (G-H) and females (I-J) treated for 8 hours in BrdU after 6 days treatment in either Estradiol, 11-Ketotestosterone, Testosterone or ethanol (control). “n” represents the number of individual fish analyzed. For each graph, one-way ANOVA with Tukey’s multiple comparison test revealed significant differences (P < 0.05) when letters are different (a, b and c).

We then investigated the effect of sex steroids on gonadotrope cell proliferation. Unfortunately, we lost part of the samples during the labeling process leading to reduced number of samples in some of the groups (see “n” in Figure 2G-J). Nevertheless, steroid treatments before BrdU incubation and labelling by IF, revealed that contrary to 11-ketotestosterone (11-KT), both estradiol (E2) and testosterone (T) were able to significantly increase the number of both BrdU/hrGfpII and BrdU/Fshβ cells in male pituitaries compared to control. In females, T was able to significantly increase the number of BrdU/Fshβ and BrdU/hrGfpII cells compared to control. In contrast, other treatments did not affect the number of BrdU/hrGfpII or BrdU/Fshβ cells in females.

### Distribution of Lh and Fsh cells in the pituitary

Based on observations in the double transgenic line (*lhb*-hrGfpII/*fshb*-DsRed2), hrGfpII and DsRed2 positive cells are distributed in the median part of the pituitary in adult fish (Figure 3). This becomes even more clear when looking at the distribution in a 3D reconstituted image of the pituitary (Juveniles: Supplemental movie 1 and 2; adults: supplemental movie 3 and 4). We did not observe any difference between sex (data in males not shown), however, we could clearly see that in adults, hrGfpII cells are situated along the ventral and lateral surface of the pituitary while DsRed2 are located more internally. While hrGfpII cells to a large extent are clustered, DsRed2 cells seem more individualized and spread out. In juveniles, some of the DsRed2 cells were closer to the surface, some even touching the ventral and lateral surface of the pituitary, while this was never observed in adults where Lh cells cover the entire ventral and lateral surface.

**Figure 3:**
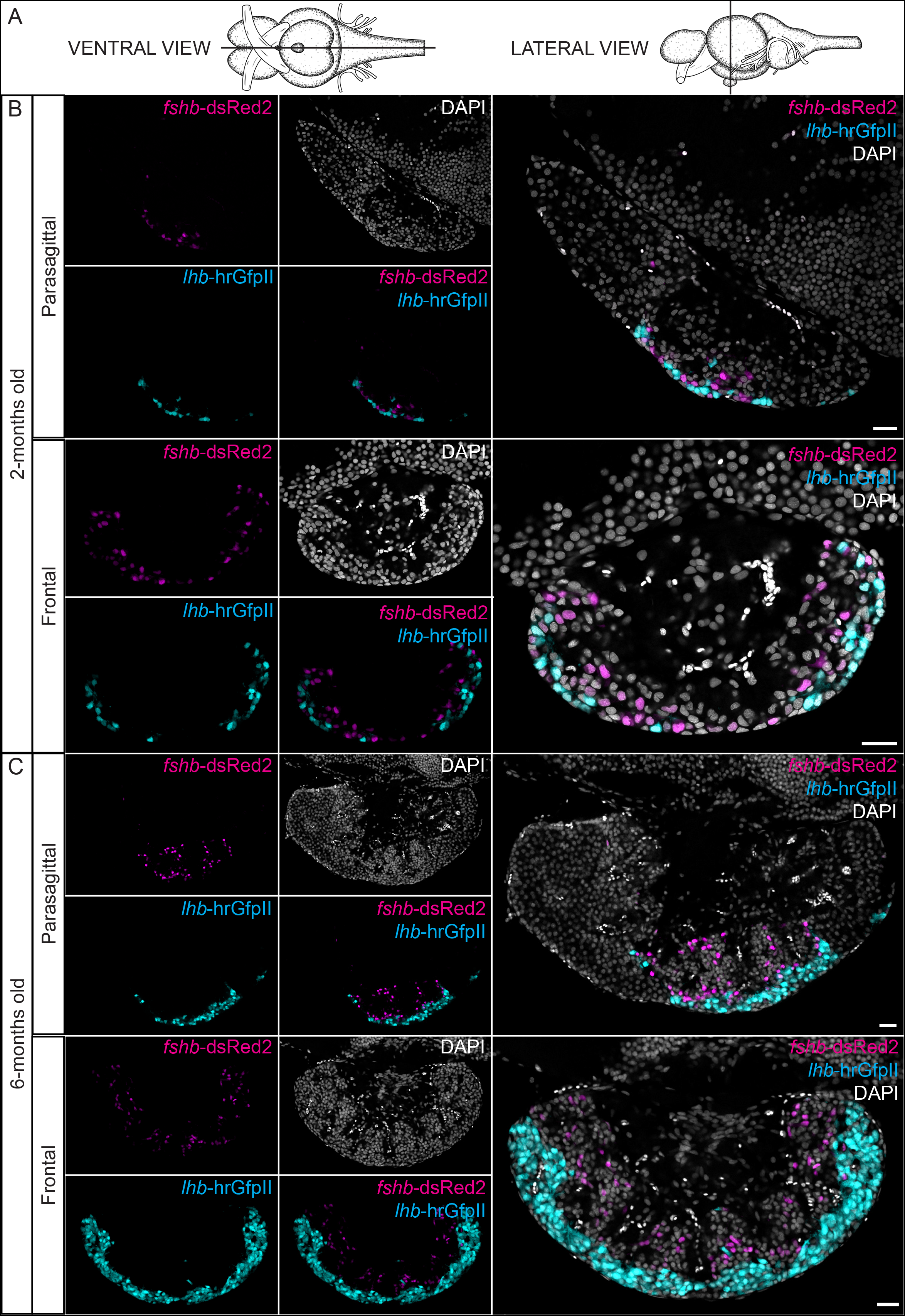
(A) Schemas presenting the position of the sections made in the brain and pituitary used for the following images, from the ventral and ventral point of view providing respectively parasagittal and frontal sections. (B) Confocal plan images of the endogenous fluorescence from 2-months old and 6-months old females tg(*lhb*-hrGfpII/*fshb*-DsRed2) medaka brain and pituitary, in parasagittal and frontal sections. Sections are shown without or with nuclear (DAPI) staining. Scale bars: 20 μm.

Interestingly, a few cells were positive for both hrGfpII and DsRed2 in 6-, 2- and 1-month old fish (Figure 4A). Such co-expression was also shown in WT animals using FISH for *lhb* and *fshb* mRNA (Figure 4B). However, cells expressing both reporter proteins were never observed in 14 dpf larvae (n=12 larvae), at which time the first Lh cells arise in the pituitary. At this developmental stage, a few cells were weakly labeled either hrGfpII or DsRed2, but never in the same cell (Figure 4A).

**Figure 4:**
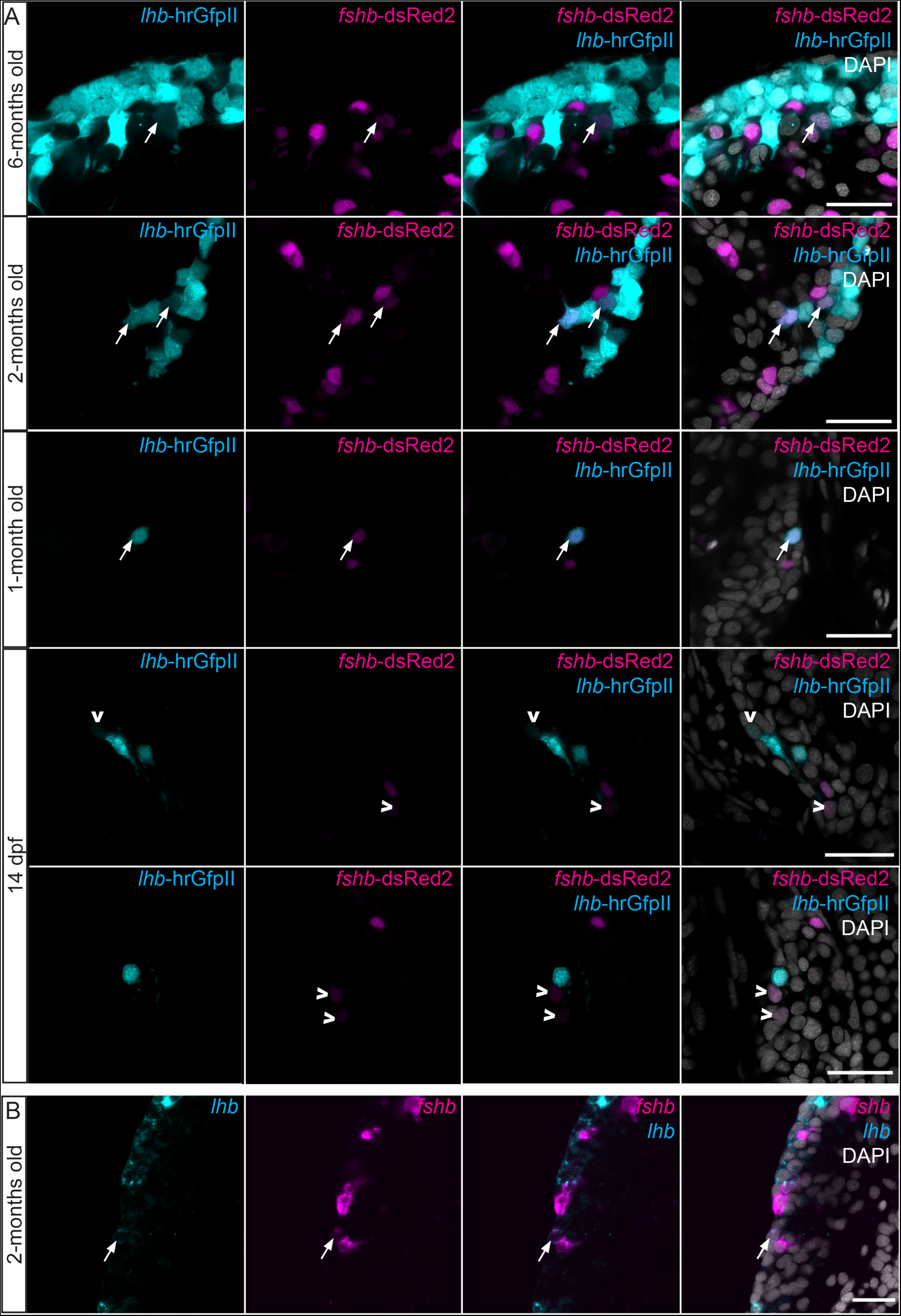
(A) Confocal plan images from the endogenous fluorescence in parasagittal sections from 6-months old, 2-months old, 1 month old and 14 dpf unsexed tg(*lhb*-hrGfpII/*fshb*-DsRed2) medaka brain and pituitary. (B) Confocal plan images of a parasagittal section from the brain and pituitary of a 2-months old WT fish labeled by multi-color FISH for *lhb* and *fshb* mRNA. Cells expressing both hrGfpII and DsRed2 (A) or *lhb* and *fshb* (B) are shown with white arrows while cells showing weak expression of DsRed2 or hrGfpII are shown with white arrow heads (A). Scale bars: 20 μm.

### Morphology of Fsh and Lh cells

Using the double transgenic line (*lhb*-hrGfpII/*fshb*-DsRed2), we investigated cell morphology. Measuring the volume of both hrGfpII and DsRed2 positive cells in dissociated pituitary cell cultures from juveniles and adults (Figure 5A), we observed a volume increase from juvenile to adult stages in both cell types, although significantly different only in adult females. Interestingly, the cell volume is similar for hrGfpII and DsRed2 positive cells at both analyzed life stages.

**Figure 5:**
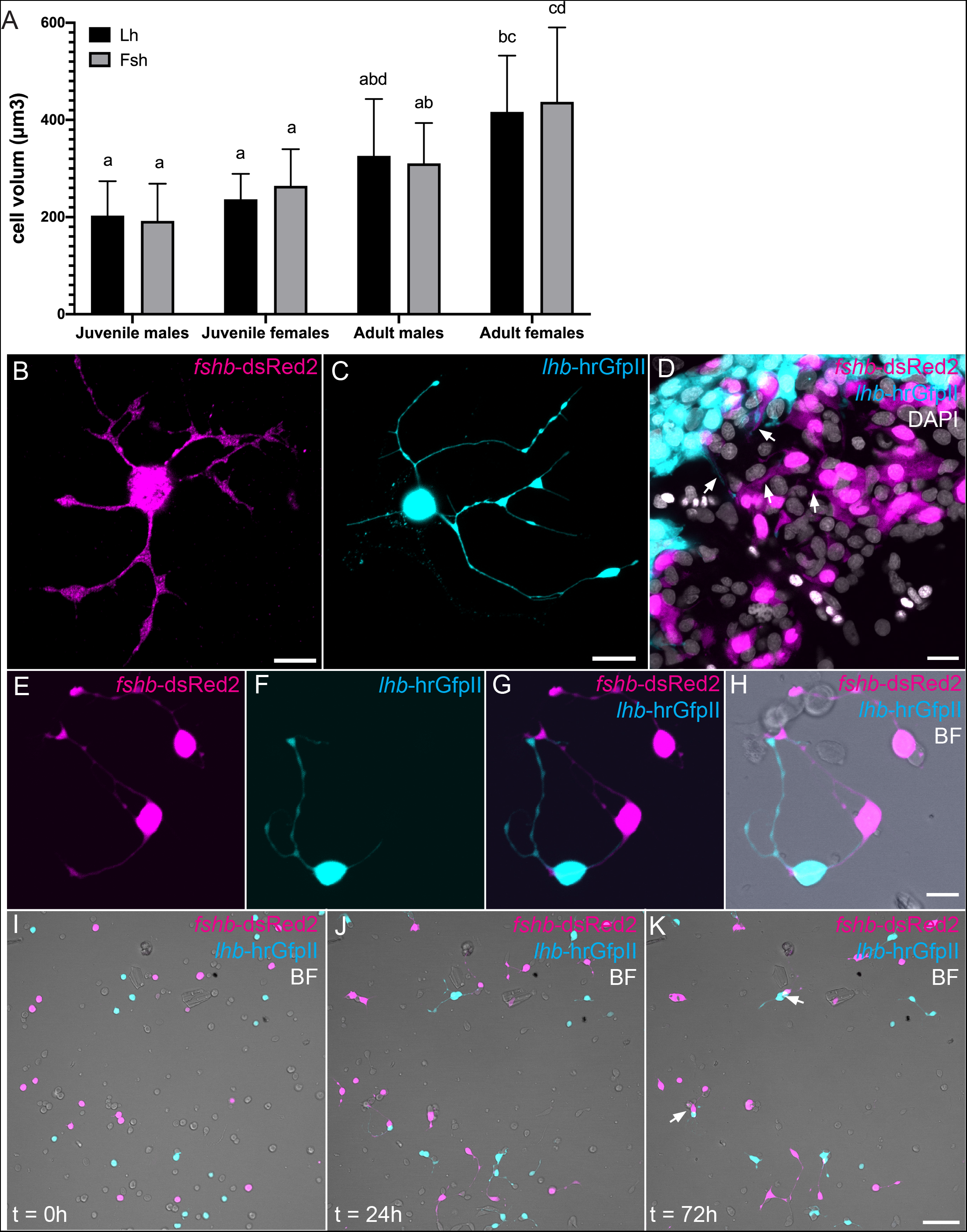
(A) Graphic showing the calculated cell volume of Lh (hrGfpII) or Fsh (DsRed2) cells in cell culture from tg(*lhb*-hrGfpII/*fshb*-DsRed2) animals, just after cells were dissociated and plated. Cell volume was measured in cells from juvenile males (n=10 cells) and females (n=9 cells) as well as in adult males (n=13 cells) and females (n=23 cells). Two-way ANOVA with Tukey’s multiple comparison test revealed significant differences (P < 0.05) when letters are different (a, b, c and d). (B,C) Confocal plan image from a dsRed positive (Fsh) and hrGfpII (Lh) cell respectively, in cell culture for 24 hours. (D) Confocal plan image from a parasagittal section of a pituitary from adult tg(*lhb*-hrGfpII/*fshb*-DsRed2) female with nuclear (DAPI) staining. Arrows show the extensions of the cells in the tissue. (E-H) Confocal plan images from pituitary cell culture from tg(*lhb*-hrGfpII/*fshb*-DsRed2) adult females, 24 hours after dissociation showing heterotypic network between a dsRed positive (Fsh) and hrGfpII (Lh) cell as well as other unknown cell types revealed by the brightfield (BF) image. Scale bars: 10 μm. (I-K) Time lapse image of a pituitary cell culture from tg(*lhb*-hrGfpII/*fshb*-DsRed2) adult females showing clustering of dsRed positive (Fsh) and hrGfpII (Lh) cells as shown by the arrows. Scale bar: 50 μm.

In addition, we observed that both in cell culture as well as in fixed tissue slices, hrGfpII and DsRed2 positive cells show seemingly similar long extensions from the cell body (Figure 5B-D). In dissociated cell culture, they use these extensions to make connections between them (homotypic and heterotypic networks; Figure 5E-H and supplemental movie 5). They also use these extensions for clustering (Figure 5I-K and supplemental movie 5).

### Phenotypic conversion of Fsh cells into Lh cells in medaka primary pituitary cell culture

Recording time lapse images of dissociated primary pituitary cell culture from the double transgenic line (*lhb*-hrGfpII/*fshb*-DsRed2), we observed that some cells which were not expressing hrGfpII at the beginning were able to start to produce it with time (Figure 6A-B and supplemental movie 6). Most of the cells starting to express hrGfpII in the culture where DsRed2 positive and some of them start to produce hrGfpII already after 15 hours in cell culture. In addition, while we observed an increase of hrGfpII fluorescence over time, we did not observe any decrease of DsRed2 fluorescence in these cells (Figure 6B). We also found that adding Gnrh1 in the medium, significantly increased the number of DsRed2 positive cells starting to produce hrGfpII (Figure 6C). Interestingly, we also observed cells that initially were not labelled starting to express hrGfpII, but we never observed any hrGfpII expressing cell starting to produce DsRed2.

**Figure 6:**
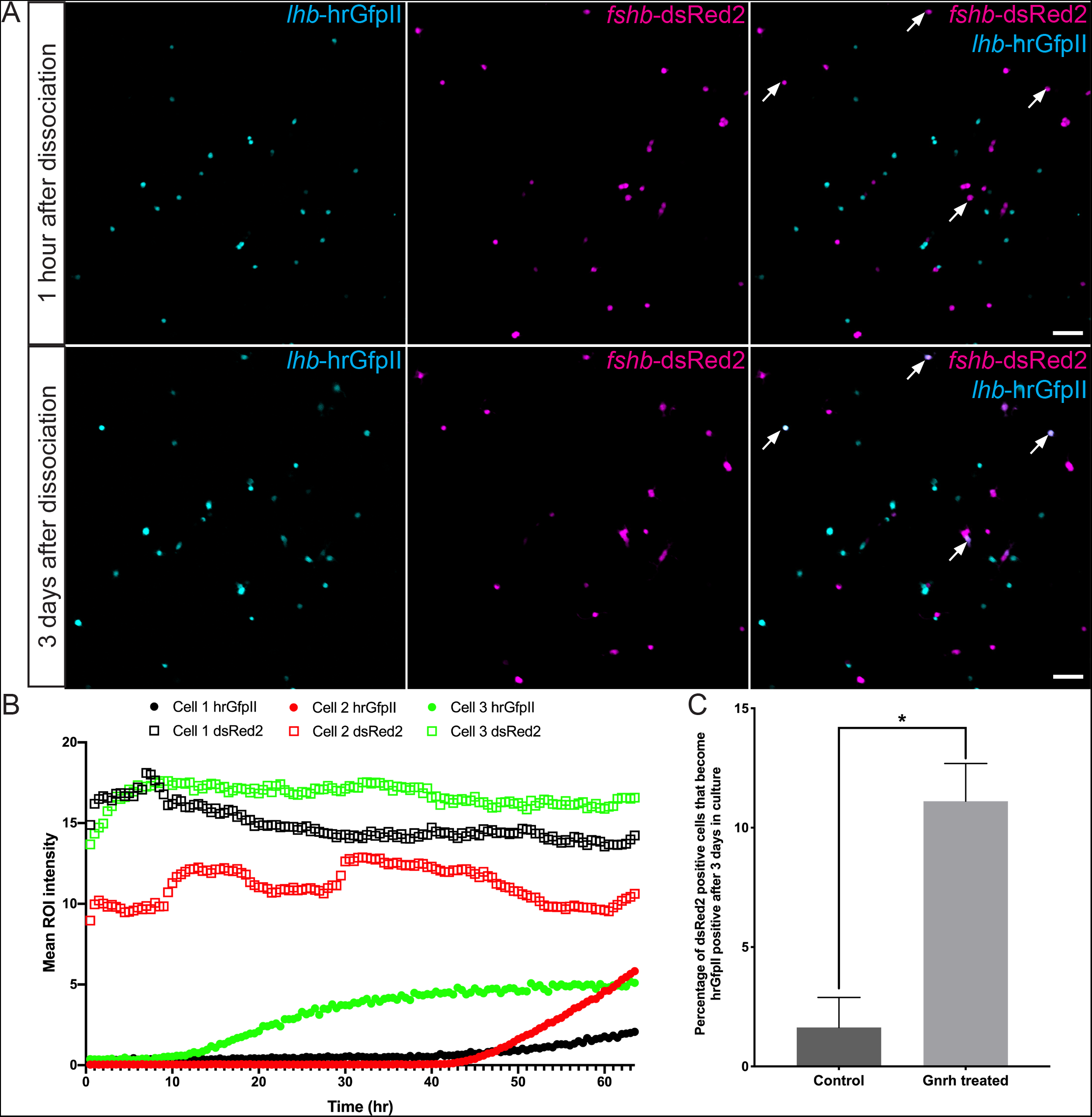
(A) Confocal plan images of a pituitary cell culture from tg(*lhb*-hrGfpII/*fshb*-DsRed2) adult males 1 hour (top panels) and 3 days (bottom panels) after dissociation. Arrows show DsRed2 positive cells that are becoming hrGfpII positive cells during the 3 days. (B) Graphic presenting the mean fluorescent ROI intensity for hrGfpII and DsRed2 from 3 different cells over time, from 2 different cell cultures imaged with a 10× objective. (C) Graphic showing the mean (+SEM) of the percentage of DsRed2 positive cells that have started to produce hrGfpII after 3 days in cell culture with Gnrh1 (n=4 cell cultures from 2 males and 2 females) or without (control n=6 cell cultures from 4 males and 2 females). Cell cultures from different sexes were pooled as they presented similar results for each treatment. Non-parametric Mann Whitney test was used to investigate significant difference in the proportion of Fsh (DsRed2) cells changing phenotype with or without Gnrh1 stimulation.

### Activity of Fsh cells upon Gnrh stimulation in medaka primary pituitary cell culture

In our previous study of adult female medaka, we demonstrated that Fsh cells lack Gnrh receptors in tissue sections and do not shown calcium or electrophysiological responses upon Gnrh stimulation when investigating the cells shortly after dissociation (Hodne, et al. 2019). Using dissociated primary pituitary cell cultures from adult female tg(*fshb*:DsRed2) line and the calcium imaging technique, we observed that 3 days after plating, 55% of the DsRed2 expressing cells show a transient elevation in cytosolic calcium with a latency of 2 to 5 sec upon Gnrh1 stimulation (Figure 7). The response usually lasted between 20 and 60 sec before returning to base line values.

**Figure 7:**
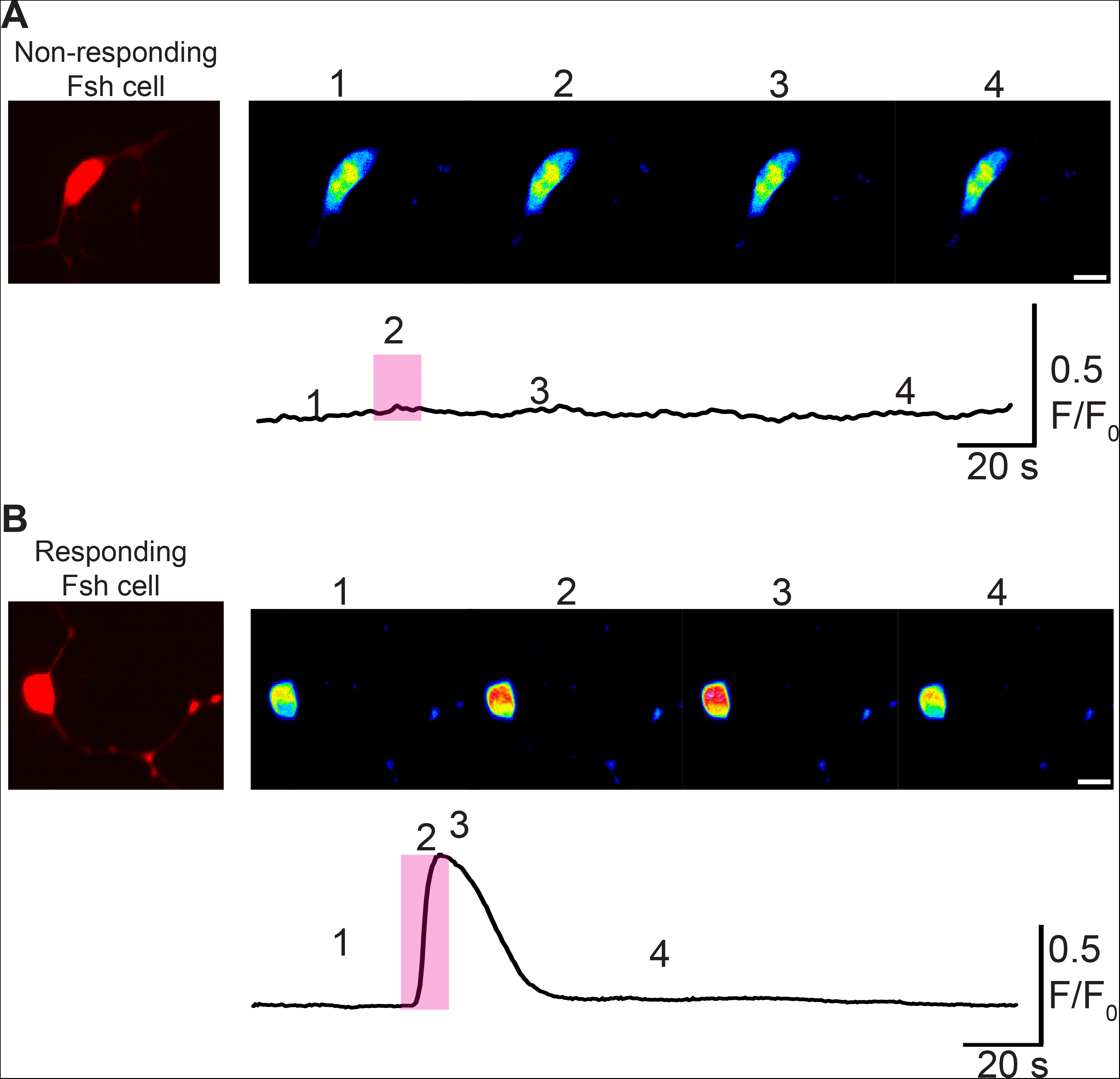
Cytosolic calcium measurements in Fsh cells following 1 μM Gnrh1 stimulation using 3 days cultivated dissociated pituitary cells from adult female tg(*fshb*-DsRed2) medaka. In total 16 of 29 Fsh cells (55%) responded to Gnrh1. Recording of the fluorescence intensity after stimulation with Gnrh1 in a (A) non-responding Fsh cell and (B) responding Fsh cell. (A and B) Upper micrographs represent four images from a time lapse of an Fsh cell following Gnrh stimulation (pink shaded rectangle). Below, The corresponding trace were each number (1-4) represents the timepoints of the selected pictures above. Scale bars on images: 10 μm.

### Temporal gene expression in primary medaka pituitary cell culture

We analyzed the gene expression over time in cell culture of *lhb*, *fshb* and the three gnrhr found in the medaka pituitary (*gnrhr1b*, *gnrhr2a* and *gnrhr2b*) according to (Hodne, et al. 2019). Three time points were studied (Figure 8), 1 hour, 24 hours and 72 hours after the dissociated cells were plated. We observed a significant reduction of the expression for *fshb* already after 24 hours. Even if not significant, we also observed a tendency to a decrease of *lhb* expression, while *gnrhr1b*, *gnrhr2a* and *gnrhr2b* expression remained stable over time.

**Figure 8:**
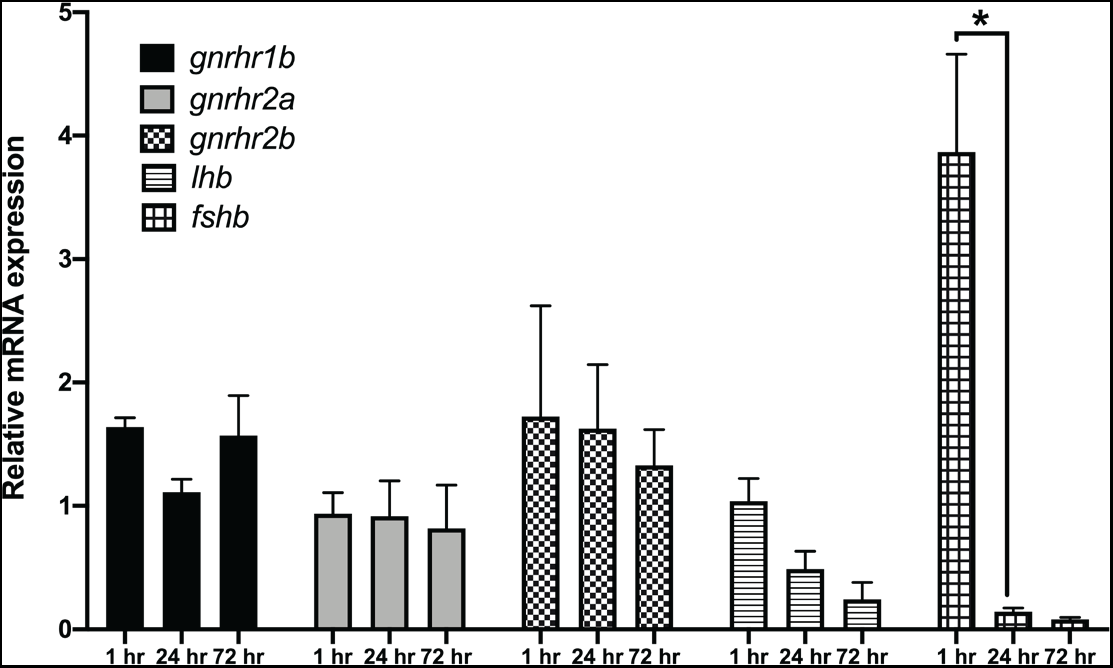
Temporal relative expression of *lhb*, *fshb*, *gnrhr1b*, *gnrhr2a* and *gnrhr2b* in cell culture from tg(*lhb*-hrGfpII) adult female pituitaries. The mRNA levels of the genes of interest were reported to the level of a combination of reference genes including *rpl7*, *gapdh* and *18s* RNA. Two-way ANOVA with Tukey’s multiple comparison test revealed significant differences (* when P < 0.05).

## DISCUSSION

Lh and Fsh are key players in the BPG axis, controlling reproductive function. While medaka Lh cells have been well described (Fontaine, et al. 2019; Hildahl, et al. 2012), little is known about Fsh cells and the population they form in the medaka pituitary. In general, less is known about Fsh cells than for Lh cells in teleost fish. In this study, we used the recently developed and validated medaka transgenic lines allowing for the visualization and localization of Fsh-producing (DsRed2) and Lh-producing (hrGfp) cells (Hodne, et al. 2019), referred to as Lh and Fsh cells, following the definition of endocrine cells used by (Pogoda and Hammerschmidt 2007).

We first studied the ontogeny of Fsh cells and demonstrated that while a significant increase of the *fshb* mRNA relative amount cannot be observed before 14 dpf, the first Fsh cell can already be observed in the pituitary after 8 dpf. This is before the observation of the first pituitary Lh cells which arise at 14 dpf (Hildahl, et al. 2012). This is comparable to zebrafish, where Fsh arise before Lh cells (respectively 4 and 28 dpf for Fsh and Lh cells; (Golan, et al. 2014)). Contrary to what has been previously described for Lh cells (Hildahl, et al. 2012), we did not observe any Fsh cell outside of the pituitary in medaka. We then observed that similar to in Lh cells (Fontaine, et al. 2019), the number of Fsh cells as well as the percentage of cells they represent in the pituitary increase between juvenile and adult stages. In addition, we demonstrate that the cell volume is also increasing between juvenile and adult stages, which is in agreement with the previous observation where Lh cell size was also observed to increase. Therefore, both the proportion and the cell volume of gonadotropes (Lh and Fsh) are increasing in the medaka pituitary between juveniles and adults in both sex, certainly because reproduction plays a more important role in adults. These observations are similar to in mammals where an increasing number and size of gonadotropes has been observed during diestrus (Childs 1986; Childs 1995). Interestingly, we noticed that the proportion of Lh cells is higher than for Fsh, in both juveniles (approx. 11% and 6%, respectively) and adults (approx. 13% and 10%, respectively).

Three hypotheses can explain the increasing number of gonadotropes in the pituitary. First, the division and differentiation of some progenitor cells. Second, the division of the gonadotrope themselves, and third, a phenotypic conversion of some of the differentiated pituitary cells. While the first hypothesis seems to have a primary role in mammalian pituitary plasticity (Florio 2011) and cannot be ruled out as some multipotent progenitor cells have been described previously in the dorsal part of the medaka pituitary (Fontaine, et al. 2019), we focused our work on the two last hypotheses.

Proliferation has previously been described for Lh cells in the medaka pituitary, and here we demonstrate that this is also the case for Fsh cells. PCNA, an essential protein for DNA replication during the cell cycle, and BrdU which has been demonstrated to be a useful and reliable marker for labelling recently divided and currently dividing cells (Bauer and Patterson, 2005), were both observed in Fsh cells, confirming active cell division. Division of hormone producing cells is not restricted to fish as this has also been observed in the mammalian pituitary (Kominami, et al. 2003) including gonadotropes themselves (Childs and Unabia 2001).

Sex steroids play crucial roles in multiple systems related to reproduction, and E2 has been shown to play an essential role in medaka reproduction (Kayo, et al. 2019). While the number of Fsh cells as well as Lh cells labeled by BrdU increased after E2 or T treatment in males, this was not the case following treatment with 11-KT (a non-aromatizable androgen). These results therefore suggest that E2, and T after aromatization into E2, are able to promote both Fsh and Lh cell proliferation in male medaka. In females however, only T was able to increase the proliferation of Fsh cells. These results are in agreement with our previous study where we observed a stimulatory effect of E2 on Lh cell proliferation in males but not in females (Fontaine, et al. 2019), perhaps due to higher endogenous levels of E2 in females (Bhatta, et al. 2012). Several studies have addressed the role of E2 and aromatizable androgens on the activity of Lh and Fsh cells in both mammals (Nett, et al. 2002) and fish (Yaron, et al. 2003). In mammals, some studies have reported a negative effects of steroids on gonadotrope cell proliferation: Mitotic gonadotropes drastically increase after castration in male rats (Sakuma, et al. 1984), and ovariectomy in female rats (Smith and Keefer 1982). In medaka however, we show a positive effect of E2 and T, on the proliferation of both gonadotrope cell types.

To test the third hypothesis about phenotypic plasticity we used the double transgenic line. We observed some gonadotropes labelled by both hrGfpII and DsRed2 in adult and juvenile stages suggesting that some cells could express both gonadotropic hormones in the medaka pituitary. We then confirmed that some cells were expressing both *lhb* and *fshb* mRNA using two colour FISH technique. Dual phenotype has been reported in other teleost fish, including the Mediterranean yellowtail (Hernandez, et al. 2002), European hake (Candelma, et al. 2017), zebrafish and tilapia (Golan, et al. 2014). It is presently unknown whether these cells are progenitor cells in a transient phenotype of differentiation toward one hormone phenotype, or fully differentiated gonadotropes in a transient form during the phenotypic conversion from one hormone phenotype to another or simply with permanent bi-hormonal phenotype. Lh and Fsh cells have been shown to share the same developmental path (Weltzien, et al. 2014), and the presence of Fsh cells has been revealed in the ventral surface of the pituitary in larval and juvenile stages, in close proximity to the Lh cells. However, we never observed dual labeling in pituitary cells of 14 dpf old larvae, the time when the first Lh cells arise. Instead, we could observe some weakly labelled hrGfpII or DsRed2 cells, suggesting that new gonadotropes arise as monohormonal cells. Therefore, the dual phenotype gonadotropes is probably not expressed in differentiating progenitor cells, but more likely in cells that change phenotype at a later stage.

We found that Lh and Fsh cells are similar in morphology. They have similar volume in juveniles and in adults and show, both in vivo and in vitro, extensions allowing networking as previously shown for Lh (Grønlien, et al. 2019). Here, we show that Lh and Fsh cells show similar behavior as they connect and cluster in cell culture using these extensions. While these similarities suggest a similar genetic background, which has already been shown (Weltzien, et al. 2014), they would also make it easy for a phenotypic conversion between the two phenotypes. We previously reported that some cells from unknown identity where able to start to produce Lh with time in cell culture (Fontaine, et al. 2019). Here, we demonstrated that in cell culture, some Fsh cells can change phenotype and start to produce *lhb*, and that Gnrh stimulates this phenotypic conversion. Interestingly, we did not observe any obvious decrease of DsRed2 fluorescence in the Fsh cells suggesting that the Fsh may become bi-hormonal, but fluorescent reporter proteins have usually relatively long half-life (about 24-30h in mammalian cells (Corish and Tyler-Smith 1999). In addition, we observed that levels of *fshb* mRNA were drastically reduced after 24 hours in cell culture, but we cannot identify which cells are responsible for this decrease. It is therefore impossible to determine if the cells that start to produce *lhb* become LH-monohormonal or bi-hormonal cells. It should also be noted that we never observed Lh cells becoming Fsh positive. These results are similar to the one observed *in vitro* in rats (Childs 1985) where mono-hormonal Fsh cells have been found to become bi-hormonal when stimulated with Gnrh. This phenotypic conversion of Fsh cells has also been described in vivo in the Rhesus Monkey during sexual maturation (Meeran, et al. 2003). In addition, it has already been described in rats (Denef, et al. 1978) and sheep (Taragnat, et al. 1998), that Gnrh was responsible for a change in the pituitary gonadotrope population by regulating the existence of LH-monohormonal, FSH-monohormonal and bi-hormonal gonadotrope subtypes. Whether this phenomenon is reversible and if other compounds could have similar or opposite effects, remains to be tested. These experiments also need to be performed ex-vivo to confirm that in medaka, phenotypic plasticity is not just due to cell culture conditions.

Interestingly, while we demonstrate here that the phenotypic conversion of Fsh cells is Gnrh sensitive, it has recently been shown that in adult female medaka, Fsh cells do not possess any *gnrhr* and do not respond (electrically nor by changes in cytosolic Ca^2+^ levels) to Gnrh stimuli after dissociation and maintained for a short period in culture (less than 48 h (Hodne, et al. 2019)). However, we show that after 3 days in cell culture, a subset of Fsh cells (about 50%) start responding to Gnrh stimuli by increasing the intracellular calcium concentration. This suggests that Fsh cells are changing phenotypic characteristics after being cultivated for an extended time without close contact with other cells, and may start to produce gnrhr. However, we did not observe any increase of *gnrhr* expression in medaka pituitary cell culture. This may be due to the relatively low number of Fsh cells in our cultures or/and the relatively high expression of gnrhr in the other pituitary cell types, thus hiding small increases of expression by Fsh cells. Indeed, a study in cod primary pituitary cell culture has reported an increase of the gene expression levels between day 2 and day 7 of one Gnrh-receptor (Gnrhr2a) found to be expressed in gonadotropes in this species (Hodne, et al. 2012). Together, these observations suggest that Fsh cells need input (paracrine or neuroendocrine factors) to maintain their status. Further studies are needed to identify the factors playing a role in the maintenance of Fsh status, but these observations further support that precautions should be taken about the conclusions when investigating dissociated primary pituitary cell cultures over time.

Many groups have reported a direct stimulation of Fsh cells by Gnrh in cell culture. While we observed that some Fsh cells start to produce *lhb* after only 15 hours in medaka cell culture, most in vitro studies use the cells several days after they were dissociated and plated: more than 3 days for coho salmon (Dickey and Swanson 2000), 5 days for rainbow trout (Vacher, et al. 2000), 2 days for masu salmon (Ando, et al. 2004), and more than 2 days for Atlantic cod (Hodne, et al. 2013). In addition, other studies have shown an effect of Gnrh on Fsh cells in more complete systems (pituitary slices or whole pituitary), where cells are kept in a more intact environment and connections with neighboring cells are preserved (tilapia (Aizen, et al. 2007) and medaka (Karigo, et al. 2014)), but a recent study showed that Fsh cells can be activated indirectly through heterotypic pituitary cell networks in medaka (Hodne, et al. 2019). Therefore, whether Gnrh directly affects Fsh cells in fish should be reinvestigated taking these new findings into account.

To conclude, this study demonstrates that gonadotropes, Lh and Fsh cells, show high plasticity by exhibiting hypertrophy and hyperplasia between juvenile and adult stages. They both proliferate in the medaka pituitary upon estradiol stimulation, and also upon testosterone stimulation after its aromatization into estradiol. Fsh cells have the capacity to change their phenotype by starting to produce Lh, and this phenomenon is promoted by Gnrh. This may explain the number of gonadotropes observed as bi-hormonal in different fish species. Combined, these two phenomena may participate in adapting hormone production to hormone demand, which differs across the life span of an animal.

## DECLARATIONS

### Ethics approval

Animal experiments were performed according to the recommendations of the care and welfare of research animals at the Norwegian University of Life Sciences, with specific approval from the Norwegian Food Safety Authority (FOTS ID 8596).

### Author’s contributions

RF, EAW, KH made the experiments. RF, EAW, KH and FAW conceived the research and analyzed the data. RF wrote the manuscript, with input from the other authors.

## Acknowledgements

We are grateful to Dr Nourizadeh-Lillabadi Rasoul for qPCR analyses, as well as Dr John Hildahl for larval sampling during early development, and Lourdes Carreon G Tan for fish facility maintenance.

---

Supplemental movie 1: 3D reconstruction of whole pituitary from tg(lhb-hrGfpII/fshb-DsRed2) juvenile female medaka imaged by LSM710 confocal with 40X oil objective and built with 3D-viewer plugin (Fiji software). Lh cells (hrGfp-II) are cyan and Fsh cells (DsRed2) are magenta. Anterior to the top.

Supplemental movie 2: 3D reconstruction of whole pituitary from tg(*lhb*-hrGfpII/*fshb*-DsRed2) juvenile female medaka imaged by LSM710 confocal with 40X oil objective and built with 3D-viewer plugin (Fiji software). Lh cells (hrGfp-II) are cyan and Fsh cells (DsRed2) are magenta. Nuclei stained with DAPI are in grey. Anterior to the top. Scale bar in red is expressed in μm.

Supplemental movie 3: 3D reconstruction of whole pituitary from tg(*lhb*-hrGfpII/*fshb*-DsRed2) juvenile female medaka imaged by LSM710 confocal with 25X oil objective and built with 3D-viewer plugin (Fiji software). Lh cells (hrGfp-II) are cyan and Fsh cells (DsRed2) are magenta. Anterior to the top. Scale bar in red is expressed in μm.

Supplemental movie 4: 3D reconstruction of whole pituitary from tg(*lhb*-hrGfpII/*fshb*-DsRed2) juvenile female medaka imaged by LSM710 confocal with 25X oil objective and built with 3D-viewer plugin (Fiji software). Lh cells (hrGfp-II) are cyan and Fsh cells (DsRed2) are magenta. Nuclei stained with DAPI are in grey. Anterior to the top. Scale bar in red is expressed in μm.

Supplemental movie 5: Confocal time-lapse recording of primary pituitary cell culture from tg(*lhb*-hrGfpII/*fshb*-DsRed2) adult male. Imaged with a LSM710 confocal and 40X oil objective in time lapse with 15 min between each picture, from 1 h after the cells have been dissociated and plated and for 72h. Lh cells (hrGfp-II) are green and Fsh cells (DsRed2) are red.

Supplemental movie 6: Confocal time-lapse recording of primary pituitary cell culture from tg(*lhb*-hrGfpII/*fshb*-DsRed2) adult male treated with Gnrh1. Imaged with a LSM710 confocal and 40X oil objective in time lapse with 15 min between each picture, from 4 h after the cells have been dissociated and plated and for 72h. Lh cells (hrGfp-II) are green and Fsh cells (DsRed2) are red.

